## Supplementary Information for "Discovering highly potent antimicrobial peptides with deep generative model HydrAMP"

### Supplementary Text

### S1 Analogue generation algorithms

#### S1.1 Baseline discovery

Supplementary Algorithm S1 was used for analogue generation (generation of active peptides similar to prototype peptide $p_{proto}$) with the baseline discovery criteria. An analogue meets the baseline discovery criteria with $\mathbb{P}_{M_{AMP}}(p) \geq 0.8$ and $\mathbb{P}_{M_{MIC}}(p) > 0.5$.

**Data:** A prototype peptide $p_{proto}$, creativity parameter $\tau \in (0, +\infty)$, number of tries for generation `nb_of_tries`.
**Result:** A new, active, and antimicrobial peptide similar to peptide $p_{proto}$ if generation is successful, `None` otherwise

$\mu_{q(\cdot|p_{proto})}, \sigma_{q(\cdot|p_{proto})} \leftarrow Enc(p_{proto})$;
$best\_candidate \leftarrow None$;
$best\_candidate\_mic \leftarrow 0.5$;
**for** *i in* `range(nb_of_tries)` **do**
  $z_{proto} \sim \mathcal{N}(\mu_{q(\cdot|p_{proto})}, \tau^2 \cdot \sigma_{q(\cdot|p_{proto})})$;
  $new\_candidate \sim Dec(z_{proto}, (c^{AMP} = 1, c^{MIC} = 1))$;
  **if** $\mathbb{P}_{M_{AMP}}(new\_candidate) \geq 0.8$ **and** $\mathbb{P}_{M_{MIC}}(new\_candidate) > best\_candidate\_mic$ **then**
    $best\_candidate \leftarrow new\_candidate$;
    $best\_candidate\_mic \leftarrow \mathbb{P}_{M_{MIC}}(new\_candidate)$;
  **end**
**end**
**return** best_candidate;

**Algorithm S1:** Analogue generation with the baseline discovery criteria for HydrAMP.

For Basic and PepCVAE models, the following algorithm was used for analogue generation with the baseline discovery criteria:

**Data:** A prototype peptide $p_{proto}$, a number of tries for generation `nb_of_tries`.
**Result:** A new, active, and antimicrobial peptide similar to peptide $p_{proto}$ if generation is successful, `None` otherwise

$\mu_{q(\cdot|p_{proto})}, \sigma_{q(\cdot|p_{proto})} \leftarrow Enc(p_{proto})$;
$best\_candidate \leftarrow None$;
$best\_candidate\_mic \leftarrow 0.5$;
**for** *i in* `range(nb_of_tries)` **do**
  $z_{proto} \sim \mathcal{N}(\mu_{q(\cdot|p_{proto})}, \sigma_{q(\cdot|p_{proto})})$;
  $new\_candidate = MAP\left(Dec(z_{proto}, (c^{AMP} = 1, c^{MIC} = 1))\right)$;
  **if** $\mathbb{P}_{M_{AMP}}(new\_candidate) \geq 0.8$ **and** $\mathbb{P}_{M_{MIC}}(new\_candidate) > best\_candidate\_mic$ **then**
    $best\_candidate \leftarrow new\_candidate$;
    $best\_candidate\_mic \leftarrow \mathbb{P}_{M_{MIC}}(new\_candidate)$;
  **end**
**end**
**return** best_candidate;

**Algorithm S2:** Analogue generation with the baseline discovery criteria for Basic and PepCVAE models.

In the algorithms above, *MAP* refers to a procedure that selects the amino acid (or the padding symbol) with the highest probability for each position in the peptide sequence.

#### S1.2 Improvement discovery

Supplementary Algorithm S3 was used for analogue generation (generation of active peptides similar to prototype peptide $p_{proto}$) with the improvement discovery criteria. An analogue $p$ meets the improvement discovery criteria when it has a larger $\mathbb{P}_{M_{AMP}}(p)$ and $\mathbb{P}_{M_{MIC}}(p)$ than the input peptide.

**Data:** A prototype peptide $p_{proto}$, creativity parameter $\tau \in (0, +\infty)$, number of tries for generation `nb_of_tries`.
**Result:** An active, and antimicrobial peptide similar to peptide $p_{proto}$ if generation is successful, `None` otherwise

$\mu_{q(\cdot|p_{proto})}, \sigma_{q(\cdot|p_{proto})} \leftarrow Enc(p_{proto})$;
$best\_candidate \leftarrow None$;
$best\_candidate\_mic \leftarrow \mathbb{P}_{M_{MIC}}(p_{proto})$;
$best\_candidate\_amp \leftarrow \mathbb{P}_{M_{AMP}}(p_{proto})$;
**for** *i in* `range(nb_of_tries)` **do**
  $z_{proto} \sim \mathcal{N}(\mu_{q(\cdot|p_{proto})}, \tau^2 \cdot \sigma_{q(\cdot|p_{proto})})$;
  $new\_candidate \sim Dec(z_{proto}, (c^{AMP} = 1, c^{MIC} = 1))$;
  **if** $\mathbb{P}_{M_{AMP}}(new\_candidate) \geq best\_candidate\_amp$ **and** $\mathbb{P}_{M_{MIC}}(new\_candidate) > best\_candidate\_mic$ **then**
    $best\_candidate \leftarrow new\_candidate$;
    $best\_candidate\_amp \leftarrow \mathbb{P}_{M_{AMP}}(new\_candidate)$;
    $best\_candidate\_mic \leftarrow \mathbb{P}_{M_{MIC}}(new\_candidate)$;
  **end**
**end**
**return** best_candidate;

**Algorithm S3:** Analogue generation with the improvement discovery criteria for HydrAMP.

For Basic and PepCVAE models the following algorithm was used for analogue generation with the improvement discovery criteria:

**Data:** A prototype peptide $p_{proto}$, a number of tries for generation `nb_of_tries`.
**Result:** A new active, and antimicrobial peptide similar to peptide $p_{proto}$ if generation is successful, `None` otherwise

$\mu_{q(\cdot|p_{proto})}, \sigma_{q(\cdot|p_{proto})} \leftarrow Enc(p_{proto})$;
$best\_candidate \leftarrow None$;
$best\_candidate\_mic \leftarrow \mathbb{P}_{M_{MIC}}(p_{proto})$;
$best\_candidate\_amp \leftarrow \mathbb{P}_{M_{AMP}}(p_{proto})$;
**for** *i in* `range(nb_of_tries)` **do**
    $z_{proto} \sim \mathcal{N}(\mu_{q(\cdot|p_{proto})}, \sigma_{q(\cdot|p_{proto})})$;
    $new\_candidate = MAP(Dec(z_{proto}, (c^{AMP} = 1, c^{MIC} = 1)))$;
    **if** $\mathbb{P}_{M_{AMP}}(new\_candidate) \geq best\_candidate\_amp$ **and** $\mathbb{P}_{M_{MIC}}(new\_candidate) > best\_candidate\_mic$ **then**
        $best\_candidate \leftarrow new\_candidate$;
        $best\_candidate\_amp \leftarrow \mathbb{P}_{M_{AMP}}(new\_candidate)$;
        $best\_candidate\_mic \leftarrow \mathbb{P}_{M_{MIC}}(new\_candidate)$;
    **end**
**end**
**return** best_candidate;

**Algorithm S4:** Analogue generation with the improvement discovery criteria for Basic and PepCVAE.

### S2 Unconstrained generation algorithms

The following algorithm is used for generation of active and antimicrobial peptides in an unconstrained manner for HydrAMP model. Here, $z$ is sampled from the refined prior $\hat{\mathbb{P}}_z^{agg}$:

**Data:** A number of tries for generation `nb_of_tries`.
**Result:** A new active and antimicrobial peptide sampled from $\hat{\mathbb{P}}_z^{agg}$ if generation is successful, `None` otherwise.

$z \leftarrow sample(\hat{P}_z^{agg})$;
$best\_candidate \leftarrow None$;
$best\_candidate\_mic \leftarrow 0$;
**for** $i \leftarrow$ `range(nb_of_tries)` **do**
  $new\_candidate \sim Dec(z, (c^{AMP} = 1, c^{MIC} = 1))$;
  **if** $\mathbb{P}_{M_{AMP}}(new\_candidate) > 0.8$ **and** $\mathbb{P}_{M_{MIC}}(new\_candidate) > best\_candidate\_mic$ **then**
    $best\_candidate \leftarrow new\_candidate$;
    $best\_candidate\_mic \leftarrow \mathbb{P}_{M_{MIC}}(new\_candidate)$;
  **end**
**end**
**return** $best\_candidate$;

**Algorithm S5:** Unconstrained generation in the positive mode for HydrAMP

For Basic and PepCVAE models, the following algorithm is used for generation of active and antimicrobial peptides in an unconstrained manner:

**Result:** A new active and antimicrobial peptide sampled from $\hat{\mathbb{P}}_z^{agg}$ if generation is successful, `None` otherwise.
$z \leftarrow sample(\hat{P}_z^{agg})$;
$new\_candidate = MAP\left(Dec(z, (c^{AMP} = 1, c^{MIC} = 1))\right)$;
**if** $\mathbb{P}_{M_{AMP}}(new\_candidate) > 0.8$ **then**
  **return** *new_candidate*;
**end**
**return** `None`;

**Algorithm S6:** Unconstrained generation in the positive mode for Basic and PepCVAE models

The algorithm above refers to the positive mode. In the negative mode we sample peptides with conditions $(c^{AMP} = 0, c^{MIC} = 0)$, and filter out peptides with $\mathbb{P}_{M_{AMP}}(p) > 0.2$.

### S3 Benchmarking classifiers

#### S3.1 Benchmarking classifiers for prediction of high activity against *E. coli*

We comprehensively evaluated the performance of 10 published AMP classifiers (AmpGram [1], AMPlify [2], AMP Scanner [3], STM [4], four models featured in the CAMPR3 database [5], and two models featured in the DBAASP database [6, 7]). For the evaluation, we collected a set of 887 AMPs known to be highly active against *E. coli* (positives) ATCC 25922 and 5794 inactive (negatives) (Supplementary Table S4). The threshold between active and inactive peptides was set to $MIC = 10^{1.5} \simeq 32$. Sequences sharing $\geq 80\%$ sequence identity are removed, and replaced with the representative sequences of the clusters using CD-HIT [8, 9]. All selected training sequences have length of at most 35 amino acids. Having this dedicated dataset, we tested accuracy, F1 score, true- and false positive rate (TPR, FPR) for each of the classifiers. Our analysis showed generally large FPR except the two classifiers featured in DBAASP (FPR $> 0.73$ for non-DBAASP models; see Supplementary Table S5). The low performance measures for these classifiers could result from the fact that the classifiers were originally designed to predict the binary status of a peptide being AMP, while the positives and negatives in our dataset were based on MIC measurements. Importantly, the MIC values do not always agree with the AMP status, namely a peptide which is reported as an AMP in a database is likely to have different MIC values for different pathogens and in particular may not always be active and have low MIC for *E. coli*. Such construction of the dedicated dataset, however, was critical for the needs of our preselection procedure. Indeed, our aim was to identify and preselect highly active peptides with low MIC values specifically for *E. coli*.

#### S3.2 Benchmarking classifiers for ranking peptides by high predicted activity

Next, we compared the classifiers by their ability to result in reliable ranking of top AMP candidates. To this end, for those models that return a numeric score alongside binary prediction (i.e. all except DBAASP models and the CAMPR3 neural network) we calculated receiver-operating curves (ROC) and precision-recall curves with their corresponding values of area under the curve (AUC) and average precision (AVP). Resulting curves exhibit a significant drop in precision for highest-scoring sequences (Supplementary Fig. S4). As our main goal at this stage was to minimize the chances of including non-AMPs among the highest scoring sequences, we selected 3 models with smallest initial decrease of precision (AMPlify, CAMPR3-rf and STM) to inform the final ranking step.

#### S3.3 Weighting the selected classifiers for combined peptide ranking

To optimize the ranking based on the predictions of the three selected classifiers, we needed to determine a set of weights with which the predictions of AMPlify, CAMPR3-rf and STM will be combined. To this end, we used a grid search over possible weights, and identified the combination of weights that yielded the best ranking on the evaluation set. Specifically, we found such nonnegative weights summing up to 1 that resulted in the highest true positive rate among the top 60 of candidate peptides per each prototype, ranked with the weighted average of the classifiers predictions (0.94 for AMPlify, 0.04 for CAMPR3-rf, and 0.02 for STM). Note that the relatively large differences in weight values can result not only from the outstanding performance of AMPlify compared to other classifiers, but also from the scale differences between the scores of the three classifiers.

#### S3.4 Benchmarking classifiers for prediction of toxicity

Since our intention was also to test the toxicity of the generated and selected peptides, we evaluated whether the existing toxicity classifiers were suitable to provide additional input for preselection of the generated candidates. To this end, we collected 6 published toxicity classifiers, out of which 1 shared usable code and 5 were available as a web service (Supplementary Table S1). We then constructed a dedicated toxicity dataset, by collecting 129 peptides tested as toxic (HC50 $<$ 32 µg/mL) against mammalian erythrocytes (positives) and 492 tested as non-toxic (HC50 $\geq$

32 µg/mL) (Supplementary Table 4). On this dedicated dataset, we evaluated the 4 usable classifiers, measuring their accuracy and F1 score. With maximum obtained F1 of 0.52 and accuracy of 0.74, we considered the predictive power of the evaluated toxicity classifiers too low to be reliably used for candidate peptide preselection. Thus, although we did compute and report the toxicity predictions of these classifiers for our generated candidates (Supplementary Table S8) we treated those as values to be experimentally validated rather as means for preselection.

### S4 Encoder and Decoder architectures

The Encoder model is a five layer architecture model with two outputs implemented in Keras [10]. The first layer is an one-hot encoded input in form of a sequence of amino acids for a peptide $p$. The second layer is an 100-dimensional embedding layer of each of the amino acids. It is followed by two bidirectional GRU [11] layers with 128 units. This layer outputs a single vector that is concatenation of the last states from each bidirectional runs. An output for $\mu_{q(\cdot|p)}$ is a Dense layer with 64 units and linear activation. An output for $\sigma_{q(\cdot|p)}$ is a Dense layer with 64 units and $f(x) = exp(x/2)$ as an activation function.

The Decoder model is a four layer model implemented in Keras [10]. Its first layer is an input $z$ from a 64-dim latent space. It is followed by an autoregressive GRU with 64 units. An initial hidden state of this layer is set to $z$ and initial input is set to 64-dim $\mathbf{0}$ vector. This is followed by a LSTM layer with 100 units and a dropout with 0.1 probability. The final layer is a Dense layer with 21 units (for every amino acid and an additional padding characters) followed by a Gumbel Softmax [12] activation function controlled by an additional temperature parameter.

### S5 Analogue and unconstrained training objectives of the HydrAMP model

#### S5.1 Analogue objective

Analogue objective aims at mimicking the analogue generation during the training process. We want the model to construct peptides similar to ones drawn from the data distribution $\mathbb{P}_{\mathscr{X}}$ but with different conditions. In order to achieve that, for each prototype peptide, $p_{proto} \sim \mathscr{X}$ we randomly sample the pair of conditions for its analogue $\mathbf{c}_{analog}$ from a product of two independent $Bernoulli(0.5)$ and then we maximize:

$$ANALOG = \mathbb{E}_{z \sim q(z|p_{proto})} \mathbb{E}_{p` \sim Dec(z, \mathbf{c}_{analog})} H_{\Sigma} \left( \mathbf{c}_{analog}, p` \right), \quad (1)$$

which is the expected $H_\Sigma$ of the sampled $\mathbf{c}_{analog}$ and the peptide $p'$ sampled from $Dec(z, \mathbf{c}_{analog})$. High values of this expectation imply that we can expect peptides sampled from $Dec(z, \mathbf{c}_{analog})$ to both resemble $p_{proto}$ (as we sample $z$ from its posterior) and satisfy the desired conditions $\mathbf{c}_{analog}$. We approximate the expectation w.r.t. $q(z|p_{proto})$ using the reparametrization trick and taking a single sample from the variational posterior distribution. Additionally, we approximate expectation w.r.t. to a sequence of categorical distributions $Dec(z, \mathbf{c}_{analog})$ using a single sample from Gumbel Softmax [12].

#### S5.2 Unconstrained objective

Unconstrained objective mimics the *de novo* generation process during the training process. As it is crucial that our model produces peptides with a provided condition, we optimize HydrAMP to preserve the random pair of conditions $\mathbf{c}_{unc}$ by a peptide generated for a random $z_{unc} \sim P_z$. A random pair of conditions $\mathbf{c}_{unc}$ is sampled from a product of two independent $Bernoulli(0.5)$ distributions. The class code preservation is obtained by maximizing the following expectation:

$$UNC = \mathbb{E}_{z_{unc} \sim \mathbb{P}_z} \mathbb{E}_{p` \sim Dec(z_{unc}, \mathbf{c}_{unc})} H_\Sigma \left( \mathbf{c}_{unc}, p` \right), \tag{2}$$

which is the sum of expected cross-entropies between sampled $\mathbf{c}_{unc}$ and probabilities of peptide sampled from $Dec(z_{unc}, \mathbf{c}_{unc})$ being antimicrobial and active. High values of this expectation imply that we can expect peptides sampled from $Dec(z_{unc}, \mathbf{c}_{unc})$ to have the desired pair of conditions $\mathbf{c}_{unc}$. We approximate expectation w.r.t. to a sequence of categorical distributions $Dec(z_{unc}, \mathbf{c}_{unc})$ using a single sample from Gumbel Softmax [12].

### S6 Extension of training objectives with regularization terms

#### S6.1 Jacobian disentanglement regularization

##### S6.1.1 Extension of the unconstrained objective with Jacobian disentanglement regularization

In order to apply the Jacobian latent regularization to the unconstrained objective, we extended it with the following term:

$$JDR_{unc} = \mathbb{E}_{z \sim P_z} JDR^{Dec}(z, \mathbf{c}_{unc}), \tag{3}$$

where $\mathbf{c}_{unc}$ is a random pair of conditions sampled the same as in case of unconstrained objective. We approximate this expectation with a single sample from $P_z$.

##### S6.1.2 Extension of the analogue mode with Jacobian disentanglement regularization

In order to apply the Jacobian disentanglement regularization to the analogue objective we extended it with the following term:

$$JDR_{analog} = \mathbb{E}_{z \sim q(z|p)} JDR^{Dec}(z, \mathbf{c}_{analog}), \tag{4}$$

where $p$ is a prototype peptide and $\mathbf{c}_{analog}$ is a random pair of conditions sampled as in a analogue objective. Once again we approximate this expectation with a single sample from $q(z|p)$ using a reparametrization trick.

#### S6.2 Latent reconstruction regularization

##### S6.2.1 Extension of the unconstrained objective with latent reconstruction regularization

In order to apply the latent reconstruction regularization to the unconstrained objective we extended it with the following term:

$$LRR_{unc} = \mathbb{E}_{z \sim P_z} \mathbb{E}_{p' \sim Dec(z, \mathbf{c}_{unc})} \left|\left| \mu_{q(\cdot|p)} - \mu_{q(\cdot|p')} \right|\right|_2^2, \tag{5}$$

where $\mathbf{c}_{unc}$ is a random pair of conditions sampled the same as in the case of unconstrained objective. We approximate this expectation with a single sample from $P_z$ and the expectation w.r.t. $Dec$ is approximated using a Gumbel Softmax [12].

##### S6.2.2 Extension of the analogue objective with latent reconstruction regularization

In order to apply the latent reconstruction regularization to the analogue objective we extended it with the following term:

$$LRR_{analog} = \mathbb{E}_{z \sim q(z|p)} \mathbb{E}_{p' \sim Dec(z, \mathbf{c}_{analog})} \left|\left| \mu_{q(\cdot|p)} - \mu_{q(\cdot|p')} \right|\right|_2^2, \tag{6}$$

where $p$ is a prototype peptide and $\mathbf{c}_{analog}$ is a random pair of conditions sampled as in a analogue objective. Once again, we approximate this expectation with a single sample from $q(z|p)$ using a reparametrization trick and the expectation w.r.t. *Dec* is approximated using a Gumbel Softmax [12].

### S7 Loss function, metrics and evaluation

#### S7.1 Loss function

##### S7.1.1 HydrAMP loss function

The total loss of the HydrAMP model is equal to:

$$\begin{aligned} LOSS = ELBO^{\beta}_{rec} + \lambda_{analog} ANALOG + \lambda_{unc} UNC + \lambda^{JDR}_{rec} JDR_{rec} + \lambda^{JDR}_{unc} JDR_{unc} + \\ \lambda^{analog}_{JDR} JDR_{analog} + \lambda^{rec}_{LRR} LRR_{rec} + \lambda^{unc}_{LRR} LRR_{unc} + \lambda^{analog}_{LRR} LRR_{analog}. \end{aligned} \tag{7}$$

where $\lambda_{analog} = 0.05$, $\lambda_{unc} = 0.05$, $\lambda^{JDR}_{rec} = 0.1$, $\lambda^{unc}_{JDR} = 0.05$, $\lambda^{analog}_{JDR} = 0.05$, $\lambda^{rec}_{LRR} = 0.1$, $\lambda^{unc}_{LRR} = 0.05$, $\lambda^{analog}_{LRR} = 0.05$.

##### S7.1.2 Basic model loss function

The optimization objective for the Basic model is the same as in Equation (7) but with a $\lambda_{unc} = \lambda^{unc}_{JDR} = \lambda^{unc}_{LRR} = \lambda_{analog} = \lambda^{analog}_{JDR} = \lambda^{analog}_{LRR} = 0.0$.

##### S7.1.3 PepCVAE model loss function

The PepCVAE objective is the same as in (7) but with $\lambda_{analog} = \lambda^{analog}_{JDR} = \lambda^{analog}_{LRR} = 0.0$.

#### S7.2 Metrics

A part of optimizing the reconstruction objective is to make the *Dec* output distribution to be likely to reconstruct the original peptide. In order to measure this qualitatively we introduced two metrics: *LengthAcc* and *AminoAcc*. We predict the beginning of the padding as the first index at which the probability of additional padding character exceeds 0.5. *LengthAcc* measures the percentage of peptides for which the end of the sequence is predicted correctly. At each position before the predicted end of the sequence we decode the amino acid with the largest likelihood. The *AminoAcc* measures the percentage of correctly decoded amino acids up to a position of a predicted beginning of the padding.

#### S7.3 Model selection

Models from each epoch are challenged with the task of generating analogues for a validation set of both positive and negative peptides. The final models are chosen by selecting the checkpoint from an epoch with the largest sum of i) positive peptides that produced improved analogues in the improvement task and ii) negative peptides that produced analogues satisfying the acceptance criteria of the discovery task, out of those epochs for which model achieved $>$ 95% of average *AminoAcc* and $>$ 99% of *LengthAcc*.

[12] Eric Jang, Shixiang Gu, and Ben Poole. *Categorical Reparameterization with Gumbel-Softmax*. 2017. arXiv: `1611.01144 [stat.ML]`.

### Supplementary Tables

| | F1-score | | Accuracy | | Sensitivity | | Specificity | |
|---|---|---|---|---|---|---|---|---|
| | Val | Test | Val | Test | Val | Test | Val | Test |
| AMP classifier | 0.868 | 0.874 | 0.868 | 0.88 | 0.831 | 0.833 | 0.904 | 0.923 |
| MIC classifier | 0.793 | 0.804 | 0.935 | 0.942 | 0.836 | 0.852 | 0.953 | 0.956 |

**Table S1** **$M_{AMP}$ and $M_{MIC}$ classifiers (collectively denoted the Classifier) metrics.** Validation score was averaged out over splits.

| Benchmark type | Class | Data source | Number of sequences |
|---|---|---|---|
| Activity | Positive (active) | DBAASP | 1023 |
| | Negative (inactive) | Uniprot | 5604 |
| | | In-house | 54 |
| Toxicity | Positive (toxic) | DBAASP | 129 |
| | Negative (nontoxic) | DBAASP | 492 |

**Table S2 Summary of dedicated benchmark datasets for activity against *E. coli* and toxicity against human erythrocytes.** For the activity benchmark, positive (active) examples are peptides selected from DBAASP with experimentally verified $MIC \leq 10^{1.5} \simeq 32$ against *E. coli* ATCC 25922; negative (assumed inactive) examples are peptides extracted from UniProt manually using search filters requiring subcellular location: cytoplasm, and excluding the following properties: antimicrobial, antibiotic, antiviral, antifungal, effector, excreted. We also included sequences from a small in-house dataset of experimentally confirmed inactive peptides. For the toxicity benchmark, positive examples are peptides selected from DBAASP with experimentally verified HC50 $< 10^{1.5} \sim 32$ µg/mL against human erythrocytes; negative examples have experimentally verified HC50 below that threshold. The datasets are shared as part of a github repository `https://github.com/szczurek-lab/hydramp`.

| | F1-score | Accuracy | True positive rate | False positive rate |
|---|---|---|---|---|
| AMPGram | 0.779 | 0.650 | 0.936 | 0.898 |
| AMPlify | 0.793 | 0.668 | 0.968 | 0.906 |
| AMP Scanner | 0.793 | 0.670 | 0.965 | 0.897 |
| CAMPR3-ann | 0.766 | 0.638 | 0.904 | 0.873 |
| CAMPR3-da | 0.790 | 0.664 | 0.961 | 0.904 |
| CAMPR3-ann | 0.789 | 0.665 | 0.952 | 0.886 |
| CAMPR3-svm | 0.773 | 0.648 | 0.911 | 0.857 |
| DBAASP | 0.750 | 0.694 | 0.701 | 0.319 |
| DBAASP genome | 0.790 | 0.742 | 0.738 | 0.251 |
| STM | 0.695 | 0.576 | 0.736 | 0.732 |

**Table S3 Evaluation results of auxiliary AMP classifiers on the dedicated dataset.**

| | $\tau$ | Generated candidates | Filtering stage I | Filtering stage II | Selected for MD simulations | Selected for experimental validation |
|---|---|---|---|---|---|---|
| Syphaxin | 1 | 47 | 3 | 3 | 3 | 0 |
| | 2 | 221 | 5 | 5 | 2 | 0 |
| | 5 | 12597 | 107 | 102 | 17 | 6 |
| OP-145 | 1 | 777 | 733 | 206 | 20 | 1 |
| | 2 | 2846 | 2604 | 736 | 19 | 2 |
| | 5 | 31070 | 24019 | 6895 | 20 | 3 |
| GQ20 | 1 | 53 | 52 | 38 | 18 | 1 |
| | 2 | 641 | 585 | 558 | 20 | 2 |
| | 5 | 23575 | 18900 | 12013 | 20 | 3 |
| Omiganan | 1 | 8 | 8 | 4 | 3 | 0 |
| | 2 | 301 | 205 | 94 | 18 | 0 |
| | 5 | 29180 | 14568 | 4488 | 20 | 6 |

**Table S4 Number of candidate sequences remaining after each stage of the preselection process.** Filtering stage I: biological filtering criteria. Filtering stage II: classifier consensus.

### Supplementary Figures

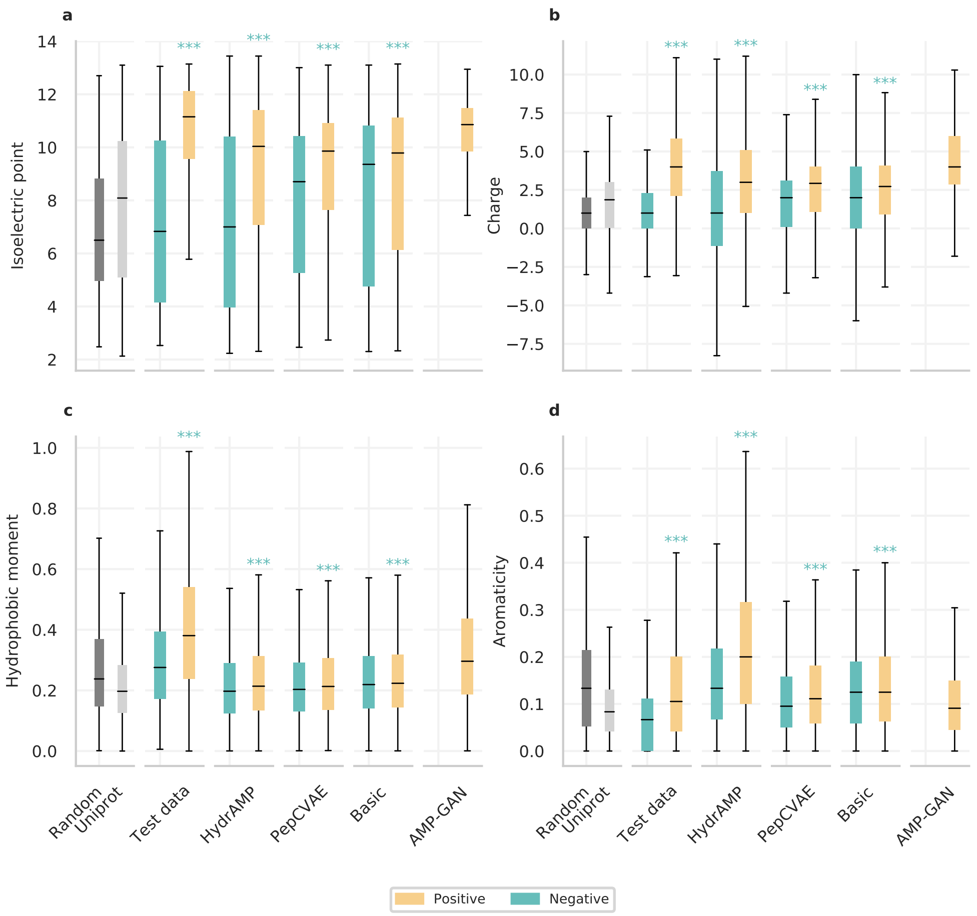


**Fig. S1 Physicochemical properties of peptides generated by HydrAMP and compared methods in unconstrained generation.** Distributions of properties of randomly generated sequences (dark gray; sample size $n = 50,000$), and peptides sampled from Uniprot (light gray; $n = 50,000$) compared with positive (yellow; $n = 50,000$) and negative (green; $n = 50,000$) peptides generated in the unconstrained generation by different models (HydrAMP, PepCVAE, Basic, AMP-GAN), and $n = 1319$ positive and $n = 1253$ negative peptides from test data. (x-axis). On y-axis: **a** Isoelectric point, **b** Charge, **c** Hydrophobic moment, **d** Aromaticity. The significance levels of one-sided Mann-Whitney test placed above the boxes are denoted as: ns - P$\geq$ 0.05; * - P$\leq$ 0.05; ** - P$\leq$ 0.01; *** - P$\leq$ 0.001. The borders of the boxes indicate first quartile (bottom) and the third quartile (top) of the data. The line within the box indicates the median. The whiskers indicate the the most extreme, non-outlier data points.

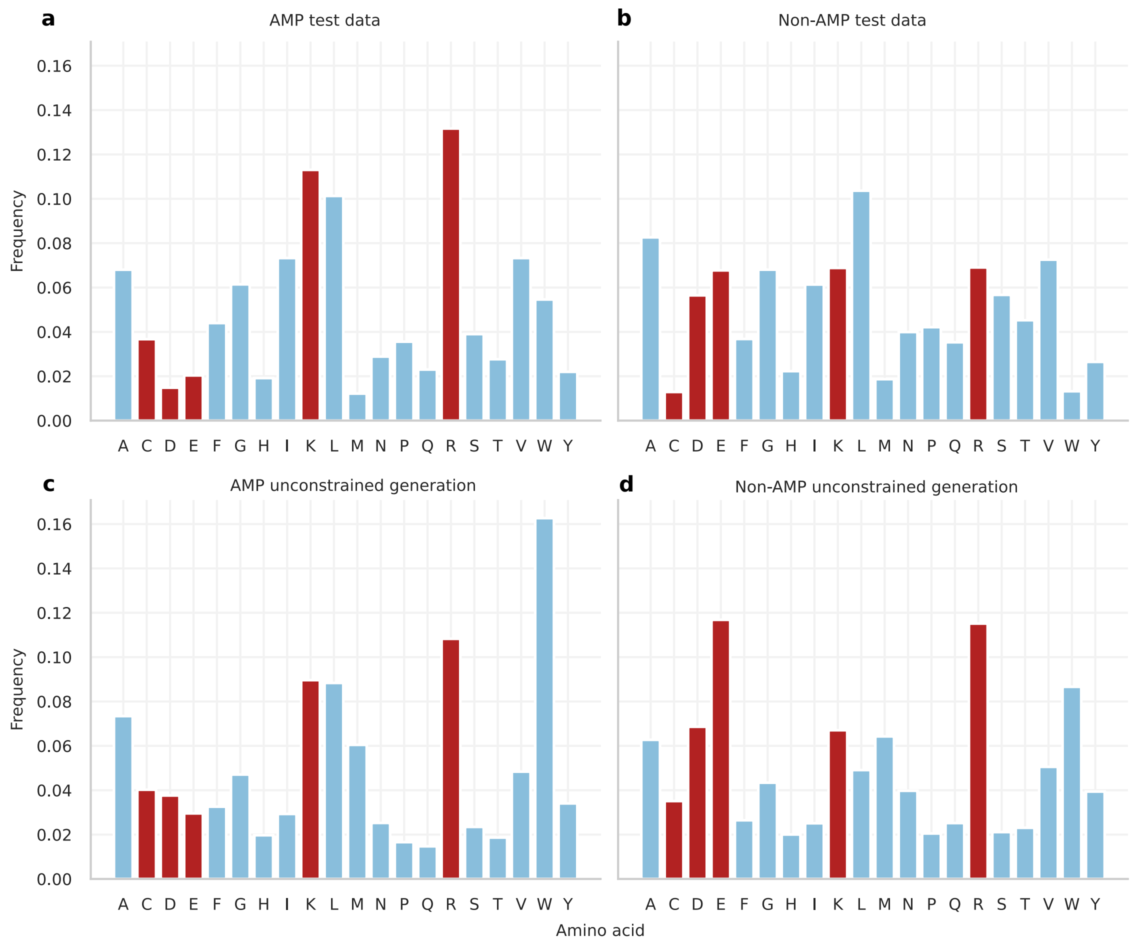


**Fig. S2 Amino acids composition of peptides generated by HydrAMP in unconstrained generation and peptides from the test set.** Bar plots of frequencies of 20 amino acids in the sequences from positive test set **(a)** and negative test set **(b)**, and in the sequences of positive **(c)** and negative **(d)** peptides generated by HydrAMP. The red bars mark amino acids contributing to charge, hydrophobicity, amphiphacity, and secondary structure.

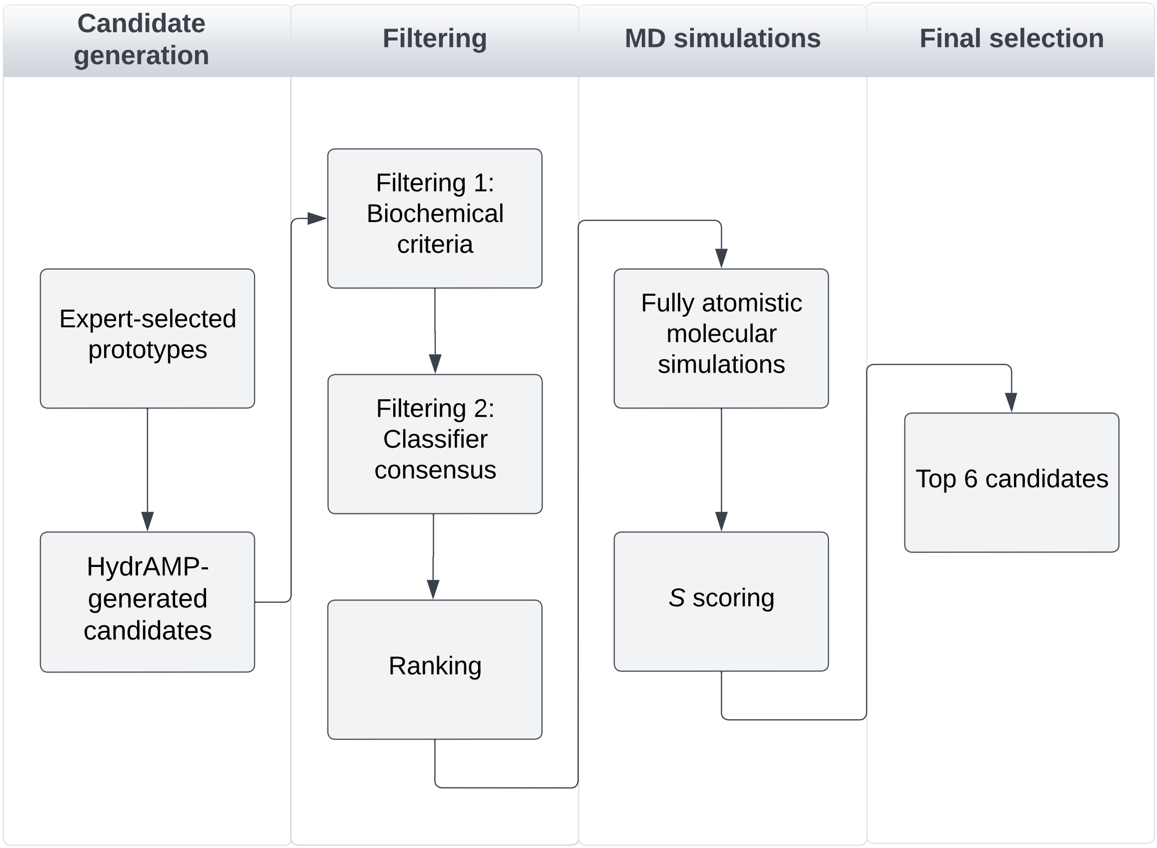


**Fig. S3 Auxiliary classifier-based ranking and simulation-based evaluation**.

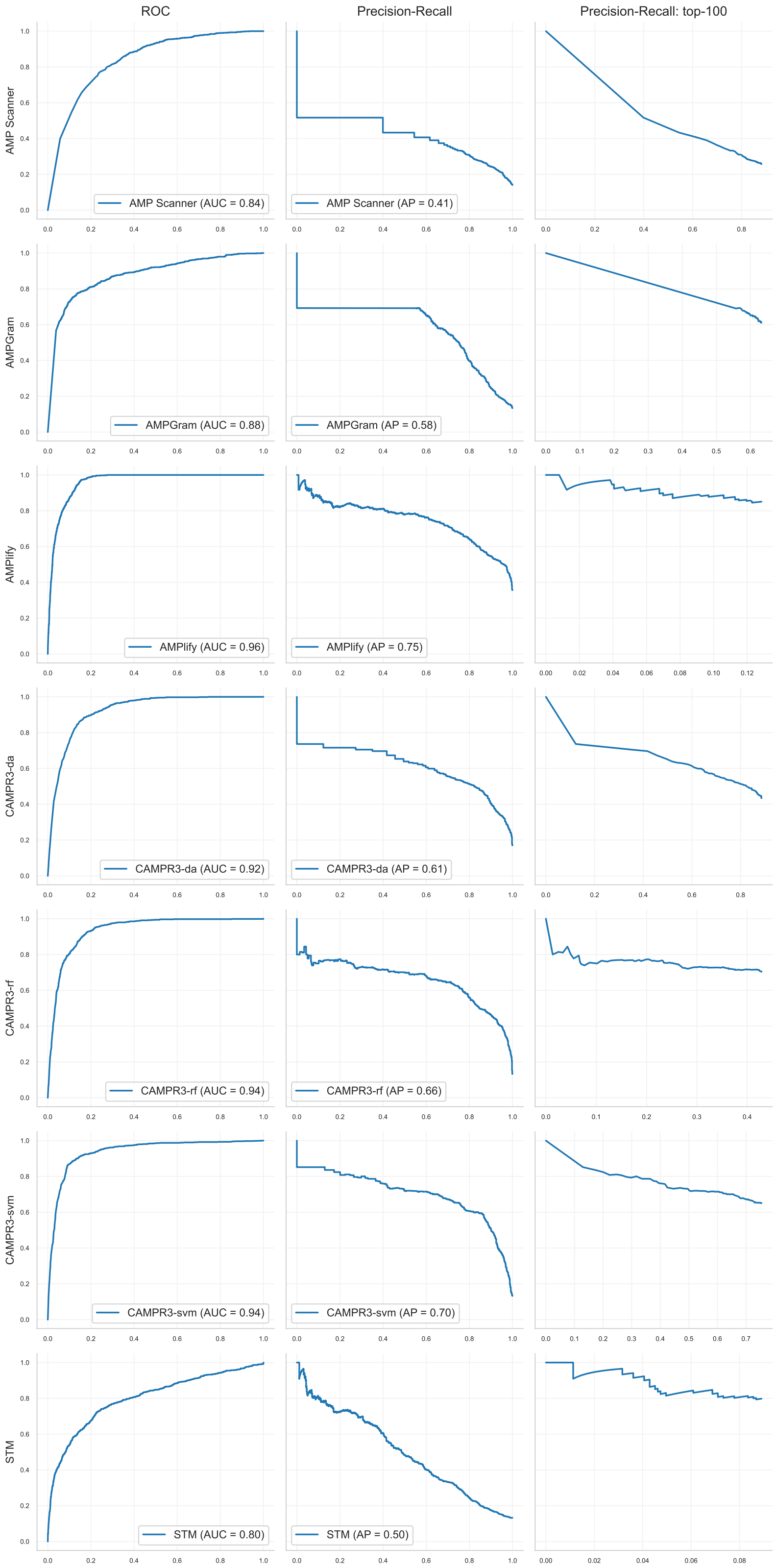


**Fig. S4 ROC, Precision-Recall and top-100 Precision-Recall curves for auxiliary AMP classifiers.**

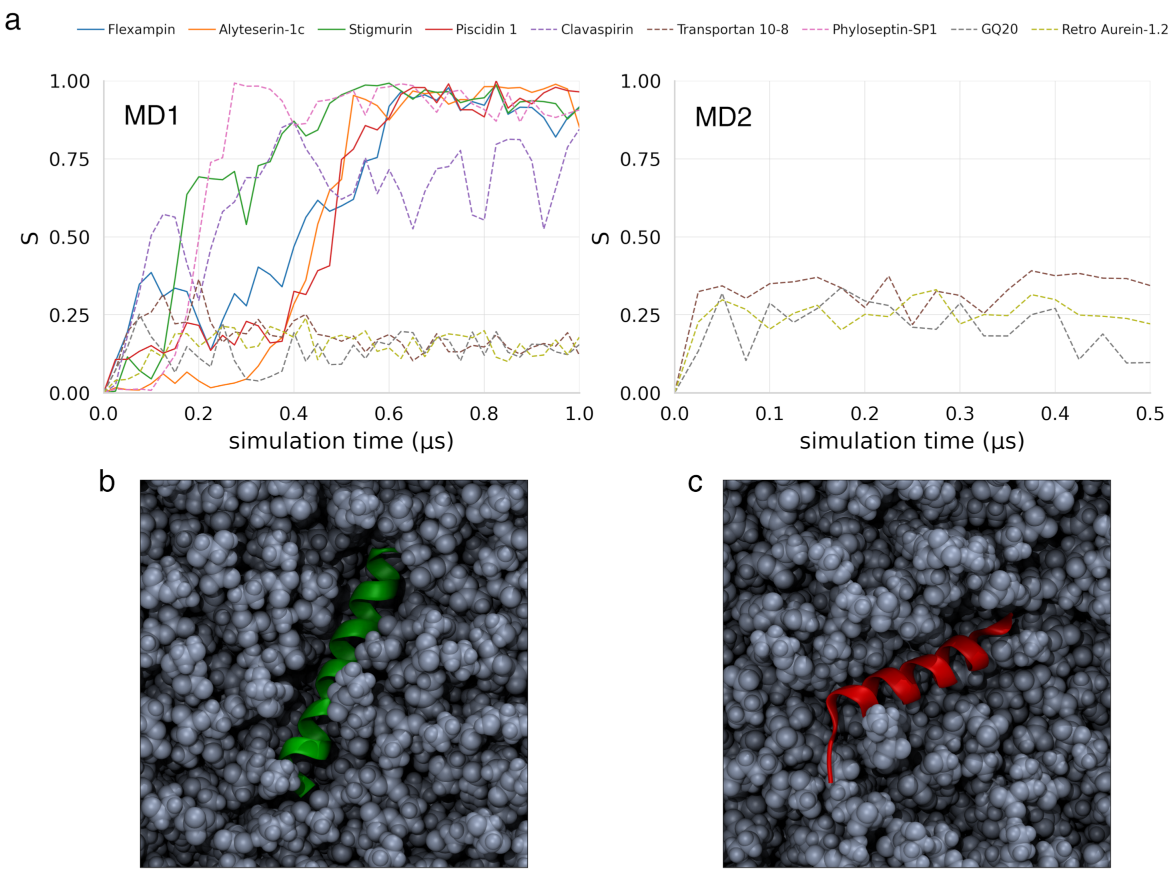


**Fig. S5 Validation of *S* parameter as AMP predictor. a** Evolution of *S* during the first (left) and the second (right) MD screening runs for experimentally confirmed active (solid lines) and inactive (dashed lines) AMPs. Note that Clavaspirin can be regarded as moderately active AMP (MIC 64 $\mu$g/mL). **b, c** Top view on membrane surface (BLKe spheres) in final simulation frames for Piscidin 1 (green) and GQ20 (red); water molecules not shown.

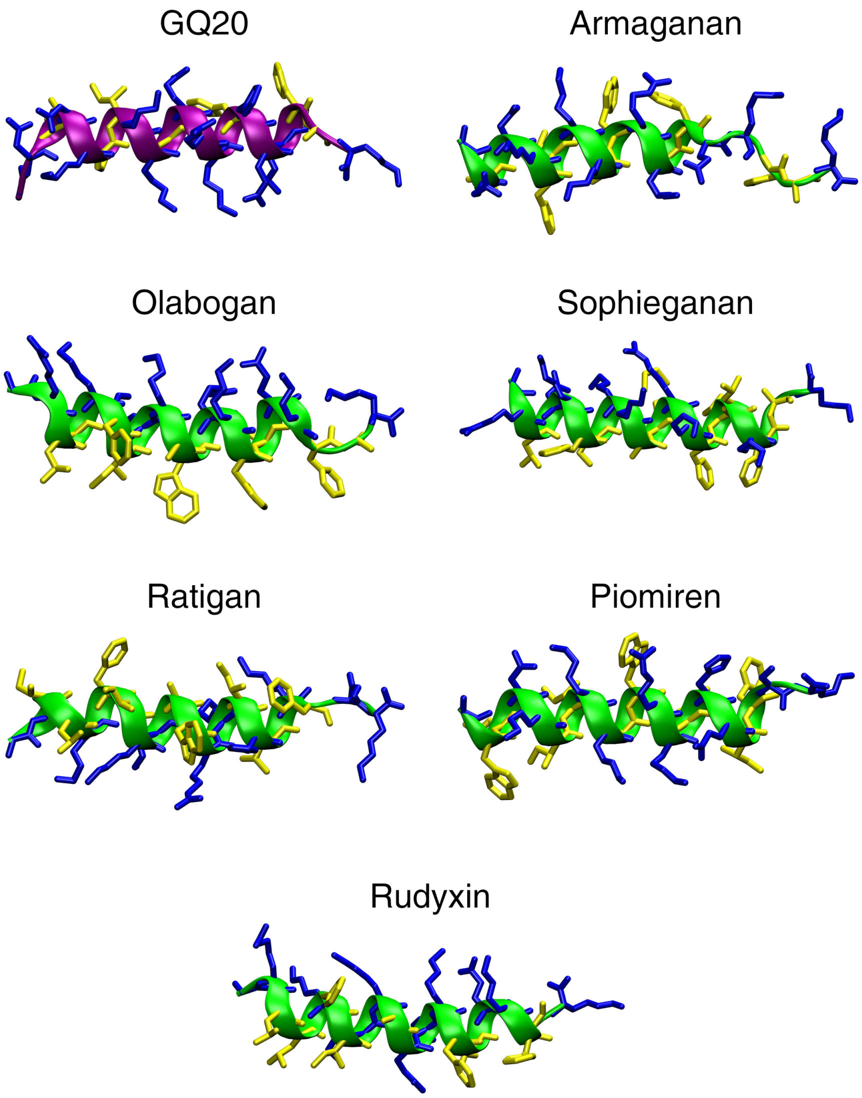


**Fig. S6 Representative structures from molecular dynamics simulations of GQ20 and its analogues.** Structures were obtained as main clusters medioids upon clustering the final 100 ns of respective runs, using gromos algorithm with 0.15 nm cutoff for heavy atom RMSD. Purple ribbon: prototype, green ribbons: active AMPs, grey ribbons: inactive AMPs. Yellow sticks: hydrophobic residues, blue sticks: polar/charged residues.

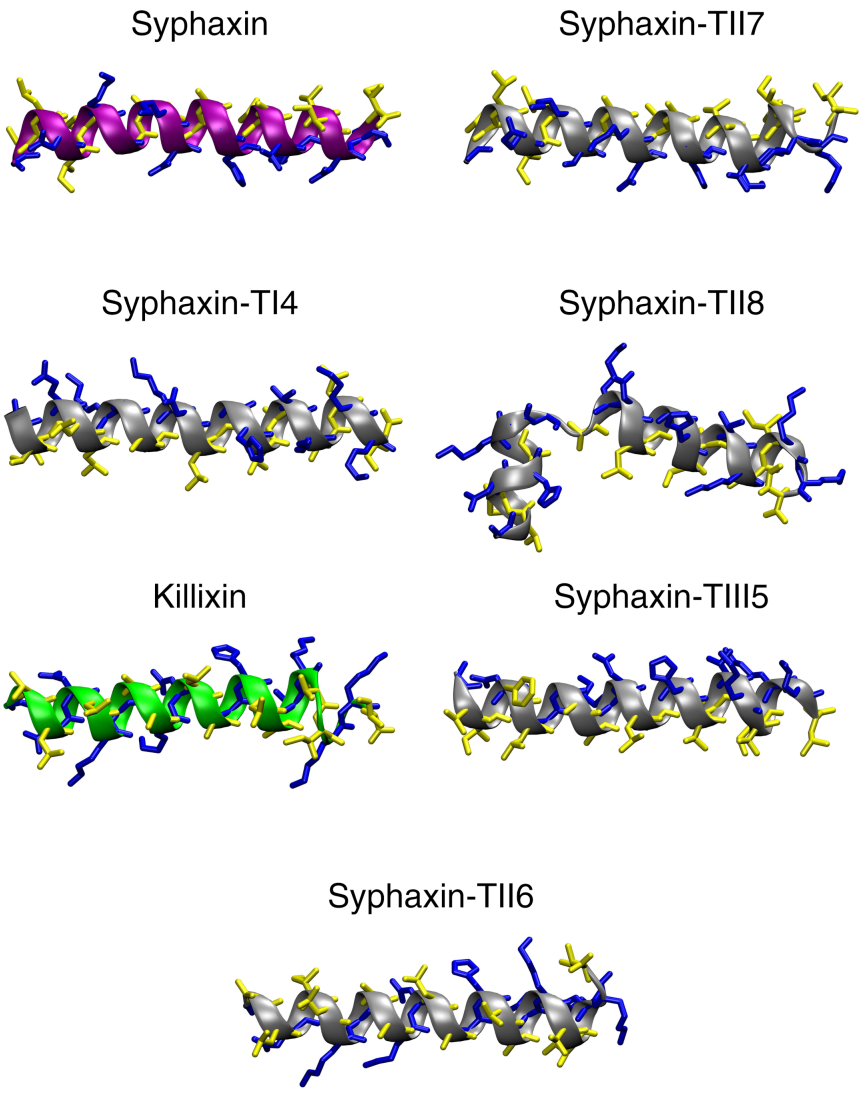


**Fig. S7 Representative structures from molecular dynamics simulations of Syphaxin and its analogues.** Structures were obtained as main clusters medioids upon clustering the final 100 ns of respective runs, using gromos algorithm with 0.15 nm cutoff for heavy atom RMSD. Purple ribbon: prototype, green ribbons: active AMPs, grey ribbons: inactive AMPs. Yellow sticks: hydrophobic residues, blue sticks: polar/charged residues.

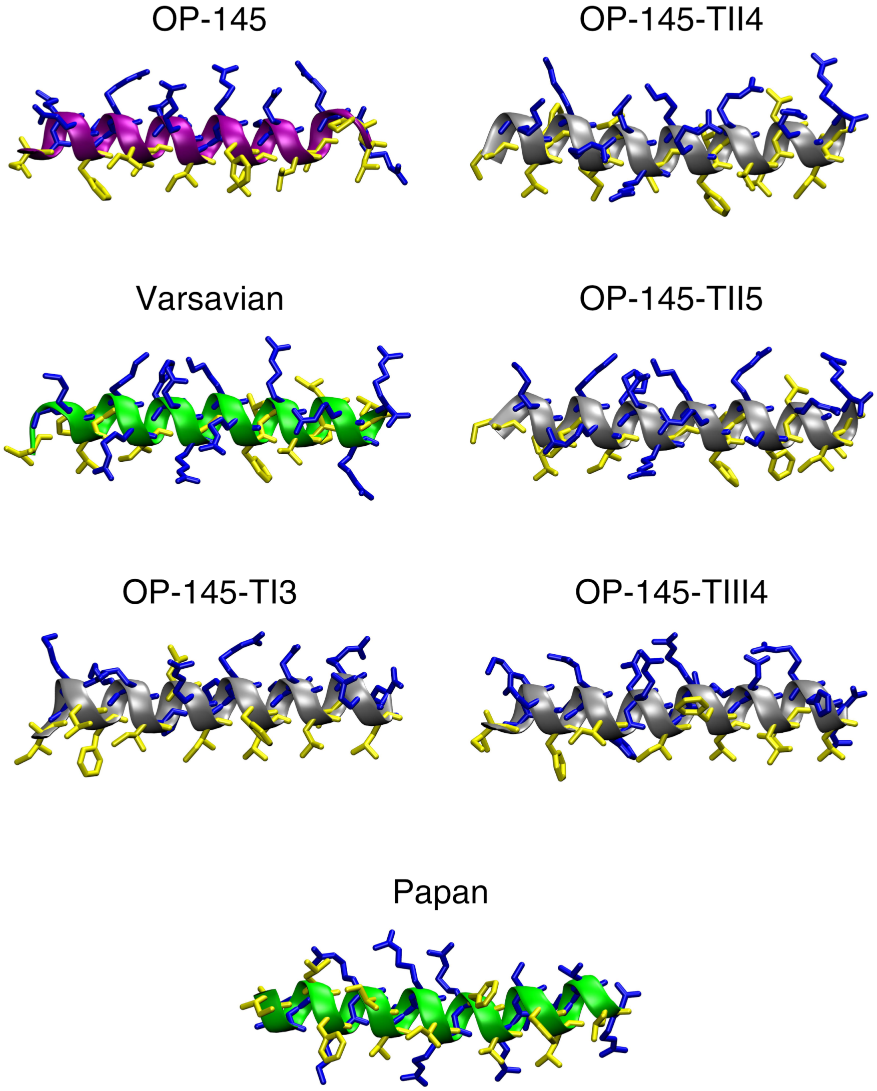


**Fig. S8 Representative structures from molecular dynamics simulations of OP-145 and its analogues.** Structures were obtained as main clusters medioids upon clustering the final 100 ns of respective runs, using gromos algorithm with 0.15 nm cutoff for heavy atom RMSD. Purple ribbon: prototype, green ribbons: active AMPs, grey ribbons: inactive AMPs. Yellow sticks: hydrophobic residues, blue sticks: polar/charged residues.

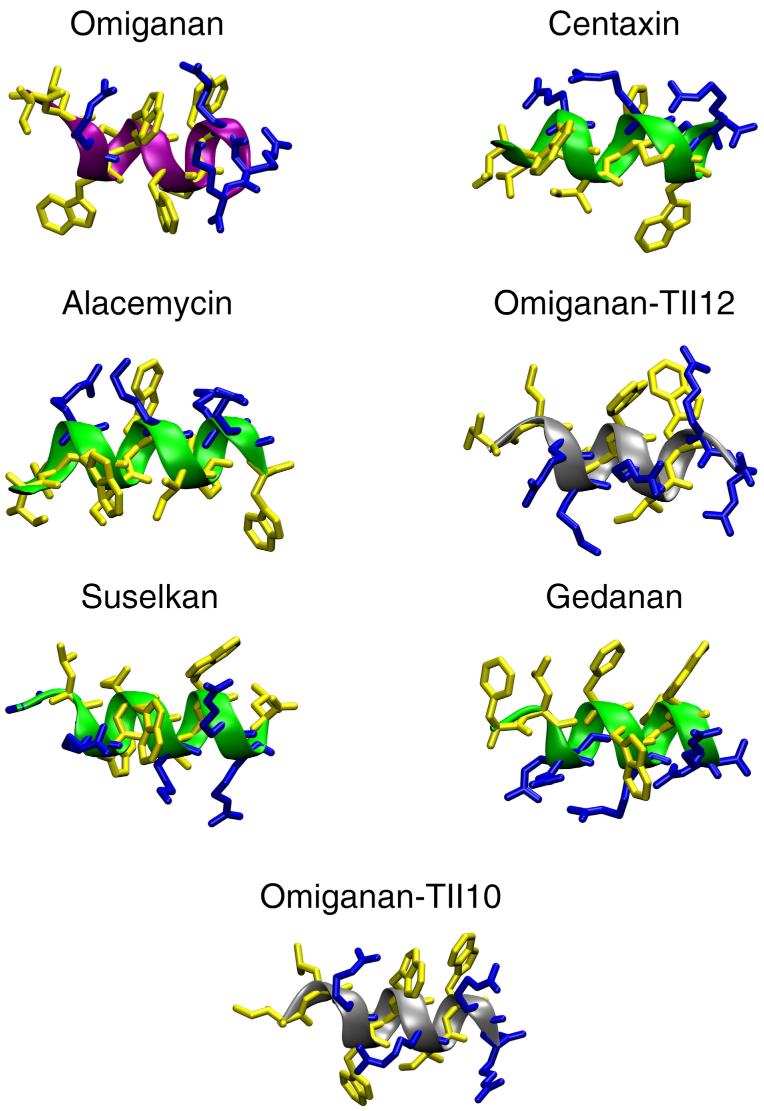


**Fig. S9 Representative structures from molecular dynamics simulations of Omiganan and its analogues.** Structures were obtained as main clusters medioids upon clustering the final 100 ns of respective runs, using gromos algorithm with 0.15 nm cutoff for heavy atom RMSD. Purple ribbon: prototype, green ribbons: active AMPs, grey ribbons: inactive AMPs. Yellow sticks: hydrophobic residues, blue sticks: polar/charged residues.

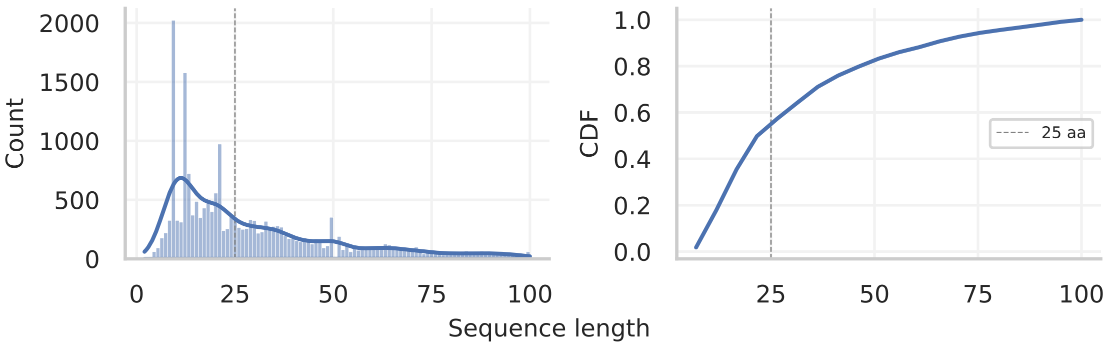


**Fig. S10 Antimicrobial peptides length distribution**. Left: Histogram of lengths of sequences shorter than 100 amino acids, right: empirical cumulative distribution function.

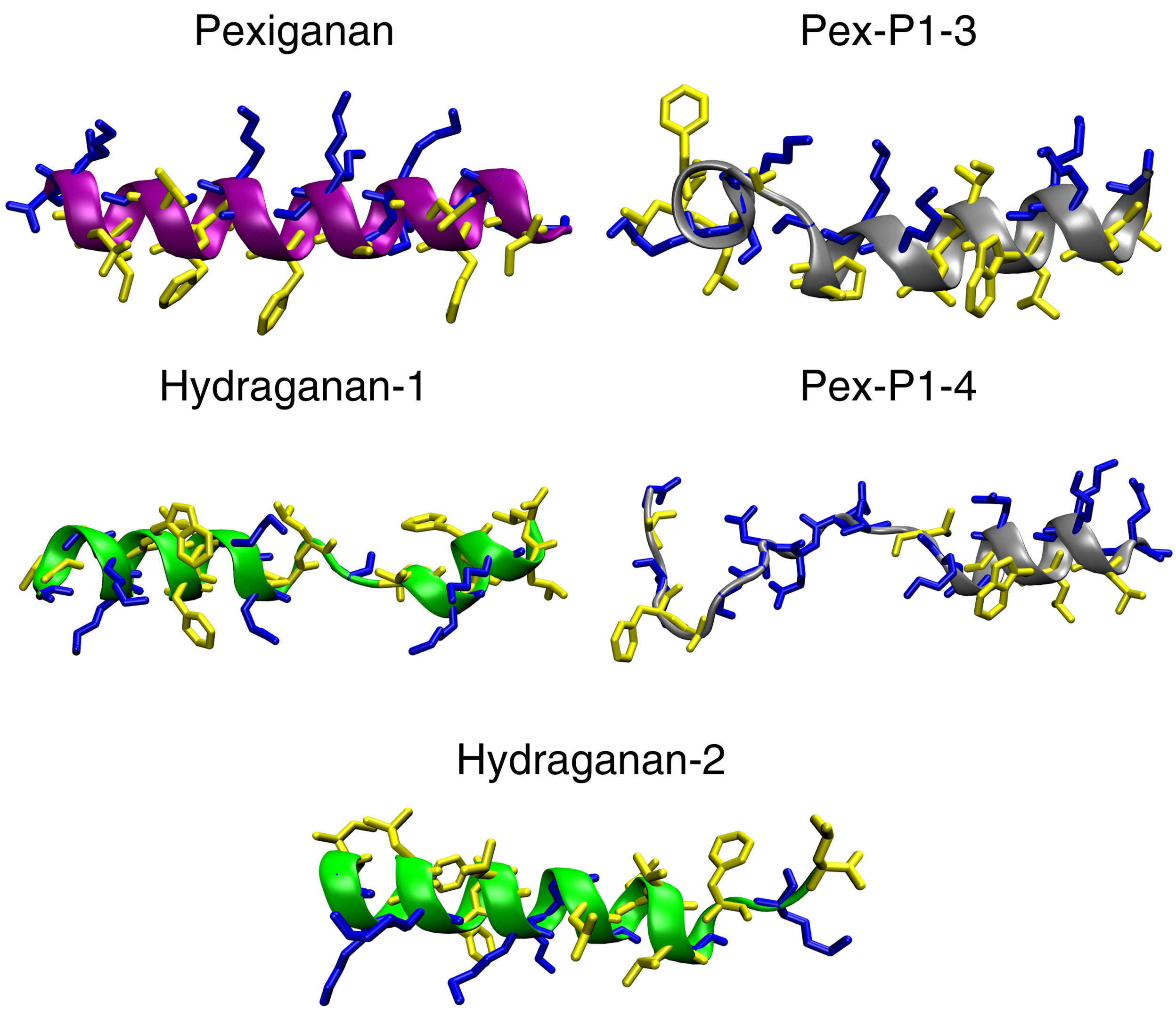


**Fig. S11 Representative structures from molecular dynamics simulations of Pexiganan and its analogues.** Structures were obtained as main clusters medioids upon clustering the final 100 ns of respective runs, using gromos algorithm with 0.15 nm cutoff for heavy atom RMSD. Purple ribbon: prototype, green ribbons: active AMPs, grey ribbons: inactive AMPs. Yellow sticks: hydrophobic residues, blue sticks: polar/charged residues.
